# Natural variation in *GmRAV* confers ecological adaptation through photoperiod control of flowering time and maturity in soybean

**DOI:** 10.1101/2021.01.11.426255

**Authors:** Yuhe Wang, Chongjing Xu, Jiafan Sun, Lidong Dong, Minmin Li, Ying Liu, Jianhui Wang, Xiaoming Zhang, Dongmei Li, Jingzhe Sun, Yuntong Zhang, Jinming Shan, Wenbin Li, Lin Zhao

## Abstract

Photoperiod strictly controlled vegetative and reproductive growth stages in soybean. A soybean *GmRAV* transcription factor containing both AP2 and B3 domains was shown to be a key component of this process. We identified six polymorphisms in *GmRAV* promoter that showed significant association with flowering time and maturity of soybean in one or multiple environments. Soybean varieties with minor polymorphism exhibited longer growth period contributing to soybean adaptation to lower latitudes. The cis-acting element GT1CONSENSUS motif of *GmRAV* promoter controlled the growth period and shortened R5-R7 by reducing the expression level of *GmRAV* in soybean. Three *GmRAV-overexpressing (GmRAV-ox)* transgenic lines displayed later flowering time and maturity, shorter height and fewer numbers of leaves compared with control plants, and transgenic inhibition of *GmRAV (GmRAV-i)* soybean displayed earlier flowering time and maturity, and increased plant heights. 163 GmRAV-target genes were determined to be putatively directly bound and transcriptionally regulated by GmRAV by combining the results from the DAP-seq and RNA-seq analyses. Two GmRAV binding motifs [C(A/G/T)A(C)ACAA(G/T)A(C/T)A(G/T)] and [C(T/A)A(C/T) C(T/G)CTG] were identified. *GmRAV* acting downstream of *E3E4* delayed soybean growth period by repressing *GmFT5a* transcriptional activity to guaranteed both vegetative and reproductive phase long enough to allow necessary energy reserved to be accumulated.

## INTRODUCTION

Photoperiod control of flowering is one of the most significant components of the interaction between plants and their environment (Borthwick and Hendricks, 1960). Different soybean cultivars exhibit different flowering and maturity patterns, which contribute to their adaptation to different ecological environments. However, as soybean is a typically photoperiod-sensitive short-day plant, the cultivation area of soybean variety is restricted to a very narrow range of latitudes (Cober and Morrison, 2010). Most soybean varieties possibly significantly reduced their yield grown in areas with 2° north latitude beyond normal cultivation latitudes (Gai and Wang, 2001). Plant adaptation to long-day conditions at higher latitudes requires early flowering and a reduction or loss of photoperiod sensitivity; adaptation to short-day conditions at lower latitudes involves delayed flowering, which prolongs vegetative growth for maximum yield potential. Therefore, soybean flowering and maturity are the key factors determining soybean productivity.

Soybean flowering time and maturity are quantitative traits controlled by multiple genes or loci. To date, forward genetic approaches have identified the major genetic loci *E1* to *E11* and *J*, and several QTLs, such as *Tof11/Gp11*, *Tof12/Gp1/qFT12-1*, and *qDTF-J*. Dominant alleles at *E1, E2, E3, E4, E7, E8*, and *E10* confer late flowering, whereas dominant alleles at *E6, E9, E11,* and *J* confer early flowering (Cober, 2011; Cober et al., 2010; Cober et al., 2001; Dissanayaka et al., 2016; Fan et al., 2014; Guo et al., 2015; Kong et al., 2010; Kong et al., 2014; Lin et al., 2020; Liu et al., 2008; Lu et al., 2017; Samanfar et al., 2016; Wang et al., 2015; Watanabe et al., 2009; Watanabe et al., 2011b; Wu et al., 2017; Xia et al., 2012; Yue et al., 2017; Zhai et al., 2014; Zhao et al., 2016). *E1* plays large effects on flowering and maturation and crucial roles in regulating photoperiod sensitivity (Han et al., 2019; Upadhyay et al., 1994; Xu et al., 2015). *E2* encodes a soybean ortholog of Arabidopsis *GIGANTEA (GmGIa)*, which plays multiple roles in the circadian clock and flowering (Watanabe et al., 2011b). *E3* (*GmphyA3*) and *E4* (*GmphyA2*) play crucial roles in regulating photoperiodic flowering under both natural daylength conditions and artificially induced LD conditions (Cober et al., 1996; Liu et al., 2008; Watanabe et al., 2009). In contrast to the *E1-E4* loci, *E6 and J* are primarily involved in promoting flowering under SDs (Bonato and Vello, 1999; Ray et al., 1995). The molecular identities of *E7* and *E8* loci are not yet known. *GmFT2a (E9*) and *GmFT5a* (*qDFT-J*) are strongly induced under SD (Kong et al., 2010) and promote flowering (Cai et al., 2018; Cai et al., 2020; Guo et al., 2015; Kong et al., 2014; Nan et al., 2014; Zhao et al., 2016). A mapping study suggests that *GmFT4* may be responsible for *E10* (Samanfar et al., 2017), which delays flowering. *E11 (Glyma.07G48500)* is one of the four soybean homologs of Arabidopsis *CCA1* and *LHY* (Wang et al., 2019). Indeed, a quadruple knockout mutant of *E11* homologs generated by CRISPR-Cas9 shows significantly delayed flowering and modified plant height (Cheng et al., 2019; Lu et al., 2020; Wang et al., 2020).

RAV subfamily members play important roles in the plant physiological processes of leaf senescence and flowering development (Zhao et al., 2017). *AtRAV2-like* and *AtRAV2* are identified as repressors of flowering in the photoperiod pathway (Castillejo and Pelaz, 2008). The presence of two DNA binding domains suggests that these RAV genes achieve high affinity by specifically binding bipartite sequences in regulatory regions of downstream targets (Castillejo and Pelaz, 2008; Kagaya et al., 1999; Matías-Hernández et al., 2016; Osnato et al., 2012). *TEM1* and *TEM2* which also belong to RAV family can delay flowering by repressing the production of FT (Matías-Hernández et al., 2014). *TEMs* also regulate GA accumulation by repressing *GA-3-OXIDASE 1 (GA3OX1)* and *GA3OX2* genes (Osnato et al., 2012). *TEM1* and *TEM2* repress trichome formation (Matías-Hernández et al., 2016), and floral transition under inductive and noninductive photoperiods(Castillejo and Pelaz, 2008; Osnato et al., 2012) as well as in response to low temperatures(Marín-González et al., 2015) and to plant age (Aguilar-Jaramillo et al., 2019). *OsRAV9/OsTEM1* has a conserved function as a repressor of photoperiodic flowering upstream of the floral activators *OsMADS14* and *Hd3a* in rice (Osnato et al., 2020).

Soybean orthologs of Arabidopsis flowering gene such as *RAV-like (RELATED TO ABI3/VP1-like)* has been characterized. *GmRAV(Glyma.10G204400)* negatively regulate SD-mediated flowering, *GmRAV*-inhibition transgenic soybean exhibited early flowering phenotype (Hu et al., 2004; Kagaya et al., 1999; Lu et al., 2014; Zhao et al., 2012). *GmRAV-03* could increase the transgenic lines resistance to high salt and drought(Zhao et al., 2017). The aim of our study is to determine the genetic effects on ecological adaptation of natural variation in GmRAV and clarify the melocular mechanism of GmRAV regulating flowering time and maturity in soybean. In this study, we detected six polymorphisms of *GmRAV* with minor allele frequencies (MAF) greater than 4% significantly or extremely significantly associated with flowering time and maturity of soybean in one or multiple environments. The cis-acting element GT1CONSENSUS motif (InDel_-904~-866normal) of *GmRAV* promoter controlled the growth period and shortened R5-R7 by reducing the expression level of *GmRAV* in soybean. *GmRAV* inhibited the expression of the *GmFT5a* by binding to CAACA and CACCTG motifs in *GmFT5a* promoter thereby delaying flowering of soybean.

## RESULTS

### Phenotypic analysis and maturity group classification

The specific photoperiod and temperature conditions resulted in distinct flowering time and maturity. Our previous work has shown that *GmRAV* negatively regulate SD-mediated flowering, *GmRAV*-inhibition transgenic soybean exhibited early flowering phenotype (Lu et al., 2014), but the genetic effect on growth periods of natural variation in *GmRAV* has not been clarified. The growth periods of 249 soybean varieties exhibited wide ranges of phenotypic variation (8.4-33.4%) which were extremely significantly influenced by genotype, environment and genotype-by-environment interaction (*P*-value<0.01) (Table S3). The *h*^2^ values higher than 59.66% reflected that the seven growth periods were primarily influenced by genetic bases (Table S3). Thirty maturity groups (MGs) standard varieties (Number 250-279) from North American were used for maturity group classification of association group in this study (Table S1), with the same planting conditions as the association group. The mid-ranges of R7 in each MGs were calculated first, and then the averages of two adjacent MGs mid-ranges were determined as the thresholds to classify neighboring MGs (Table S4) (Jia et al., 2014). The 249 soybean varieties were classified into appropriate MGs (Table S1) which were consistent with that done by Gai and Wang (Gai and Wang, 2001; Wang and Gai, 1999; Zhao et al., 2018). The MGs 000-III standard varieties matured in six environments, the MGIV standard varieties matured in three environments (2015 S, 2016 C, 2016 S), and the MGs V-VIII did not mature in any environment (Table S4).

### Polymorphisms-growth periods association analysis

103 polymorphisms comprising 90 SNPs (single nucleotide polymorphisms) and thirteen InDels (insertions and deletions) (Table S5) were identified in the entire sequence including 1,459-bp promoter, 457-bp 5’UTR and 1,056-bp CDS of *GmRAV* gene from 249 soybean varieties (Supplemental Figure 1A). Of all the 103 polymorphisms (Table S5, Table S6), 63 polymorphisms comprising 54 SNPs and nine InDels were located in the promoter region, seven polymorphisms including three SNPs and four InDels were located in the 5’UTR region, and 33 SNPs were located in the CDS region of *GmRAV*. The plot of squared allele frequency correlations (r^2^) by physical distance between pairs of polymorphisms showed a LD (linkage disequilibrium) decay distance of 1 kb (r^2^>0.1) among the entire region (2,972bp) of *GmRAV* (Supplemental Figure 1B).

To elucidate the effects of mutations of *GmRAV* on soybean growth periods, association analysis was conducted by both genotype data of 26 polymorphisms with MAF higher than 4% and phenotypic data of seven growth periods of 249 soybean varieties. Plot of the first three principal components of principal component analysis (PCA) of population structure based on genetic data obtained by SLAF-seq (Han et al., 2015) reflected a subpopulation structure of association population (Supplemental Figure 1C). Association mapping revealed that six polymorphisms (InDel_-1120T>deletion, InDel_-904normal>deletion(39bp), InDel_-655T>deletion, InDel_-557T>deletion, SNP_-293C>A, InDel_-201~-201TA>deletion) exhibited significantly (*P*-value<0.05) or extremely significant (*P*-value<0.01) association with growth periods (Supplemental Figure 1D). All soybean varieties with minor polymorphism exhibited extremely significantly longer growth periods. Soybean varieties with minor polymorphism InDel_-1120deletion exhibited longer R7 in two environments, InDel_-904deletion (39bp) showed longer R5, R6, and R7 in at least two environments, InDel_-655deletion showed longer R2, R3 and R7 in one or more environments, InDel_-557deletion showed longer R1, R2, and R7 in at least two environments, SNP_-293A showed longer R6 and R7 in one or more environments, and InDel_-201~-201deletion showed longer R1, R2, and R3 in corresponding environments (Supplemental Figure 2).

### Geographical and MGs distribution of soybeans with polymorphisms

Most of the soybean germplasms (99.6%) originated from China, therefore the distinctive geographic and MGs distribution patterns of the 248 soybean varieties from China with six polymorphisms closely related to growth periods were further estimated (Figure 1). The soybean production areas of China were divided into six ecological areas or ten sub-areas (Wang and Gai, 2002). The distributions of the soybean varieties with various minor polymorphisms were different across ecological areas (Figure 1A). Soybean varieties with InDel_-1120deletion were mainly distributed in higher latitude region in Northeast spring planting ecological sub-ecoregion, and rarely distributed in other eco-regions. Soybean varieties with InDel_-904deletion (39bp) from lower to higher latitude regions such as the eco-regions south of the Qinling Mountains-Huaihe River, Huanghuaihai double cropping planting eco-region and Northeast spring planting ecological sub-ecoregion gradually decreased. Soybean varieties with InDel_-655deletion were mainly distributed in higher latitude region in Northeast spring planting ecological sub-ecoregion, rarely distributed in the eco-regions south of the Qinling Mountains-Huaihe River and non-distributed in Huanghuaihai double cropping planting eco-region. The soybean varieties with InDel_-557deletion were mainly distributed in the eco-regions south of the Qinling Mountains-Huaihe River, less distributed in Northeast spring planting ecological sub-eco-region and non-distributed in Huanghuaihai double cropping planting eco-region. The soybean varieties with SNP_-293A were mainly distributed in the eco-regions south of the Qinling Mountains-Huaihe River, less distributed in Northeast spring planting ecological sub-ecoregion and Huanghuaihai double cropping planting eco-region. The soybean varieties with InDel_-201~-200deletion were distributed widely and regularly throughout various regions, gradually decreased from lower to higher latitude regions: the eco-regions south of the Qinling Mountains-Huaihe River, Huanghuaihai double cropping planting eco-region, and Northeast spring planting ecological sub-ecoregion. The percentages of earlier maturing varieties (MGs 000-0) with minor polymorphisms were less than those with major polymorphisms (Figure 1B).

**Figure 1.**
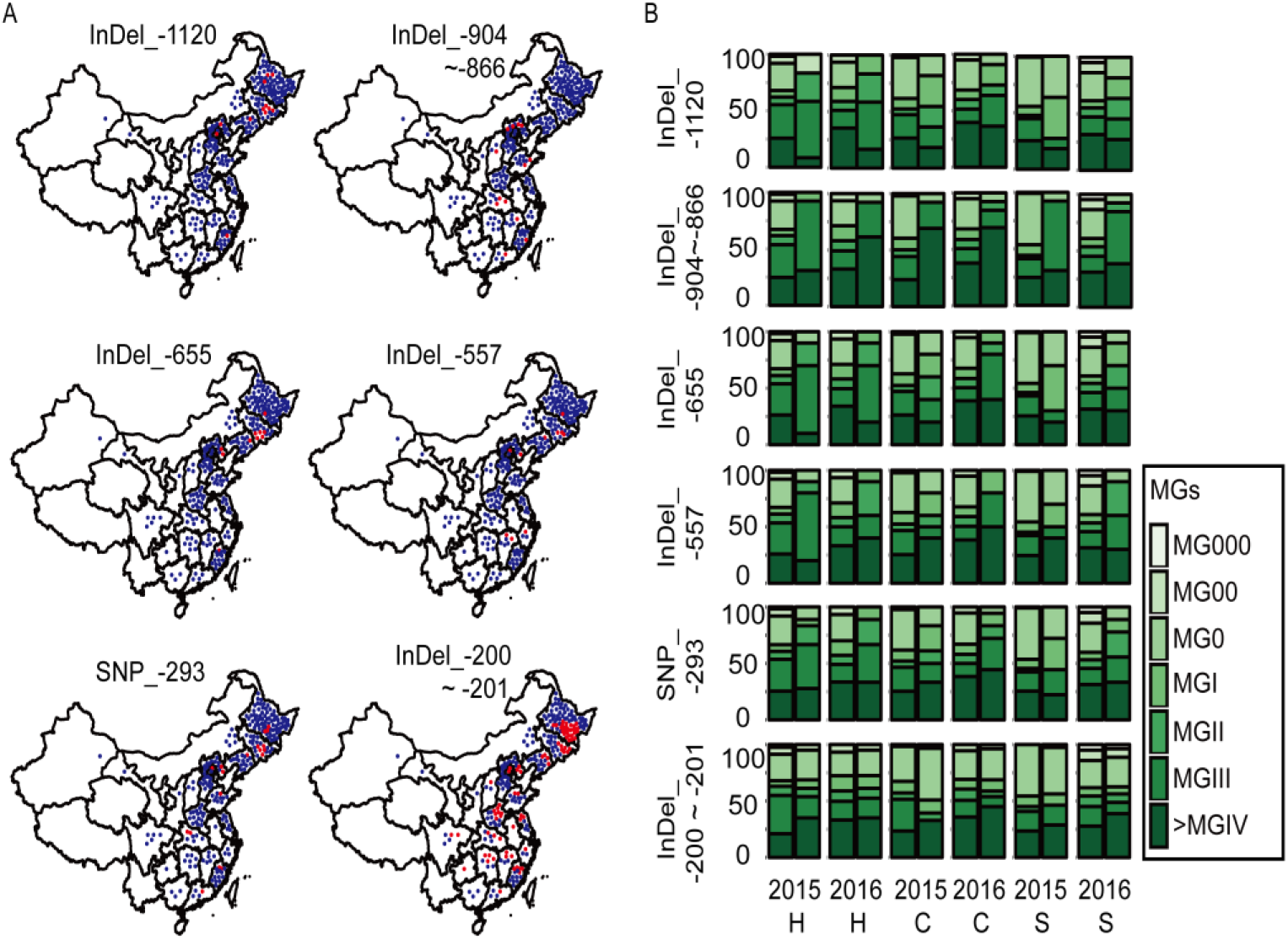
The geographical and MGs distributions of the soybean varieties with six significant polymorphisms (InDel_-1120, InDel_-904~-866, InDel_-655, InDel_-557, SNP_-293, InDel_-201~-201). (A) Geographical distribution of the soybean varieties with six polymorphisms. For each polymorphism, blue solid dots indicated the soybean varieties with major polymorphism such as InDel_-1120T, InDel_-904~-866normal, InDel_-655T, InDel_-557T, SNP_-293C, InDel_-201~-201TA, and red solid dots indicated the soybean varieties with minor polymorphism such as InDel_-1120deletion, InDel_-904~-866deletion(39bp), InDel_-655deletion, InDel_-557deletion, SNP_-293A, InDel_-201~-201deletion. (B) MGs distribution of the soybean varieties with six polymorphisms. For each polymorphism, the left column indicated the soybean varieties with major polymorphism such as InDel_-1120T, InDel_-904~-866normal, InDel_-655T, InDel_-557T, SNP_-293C, InDel_-201~-201TA, the right column represented the soybean varieties with minor polymorphism such as InDel_-1120deletion, InDel_-904~-866deletion(39bp), InDel_-655deletion, InDel_-557deletion, SNP_-293A, InDel_-201~-201deletion. X-axis showed six environments. 2015 H: 2015 Harbin; 2015 C: 2015 Changchun; 2015 S: 2015 Shenyang; 2016 H: 2016 Harbin; 2016 C: 2016 Changchun; 2016 S: 2016 Shenyang. Y-axis showed the percentages of MGs for the varieties with specific polymorphisms.

### The assays of GmRAV promoter activity and the GmRAV protein expression

Determining whether the mutations were located in known regulatory motifs such as cis-acting elements could roughly estimate whether the mutations affected the expression of functional gene. The *GmRAV* promoter sequences with polymorphisms were analyzed using PLACE programs (http://www.dna.affrc.go.jp/PLACE/) to determine the effects of polymorphisms on cis-acting elements (Higo et al., 1998). In the present study, the significant polymorphism InDel_-904deletion(39bp) (sense strand) in *GmRAV* promoter caused the variation of GT1CONSENSUS motif involved in light-responsive actions (Xue et al., 2018) (Figure 2A). The *GmRAV* promoter fragments with polymorphic InDel_-904~-866 normal and InDel_-904~-866 deletion (39bp) (Table S7), from soybean varieties ‘Dongnong 42’ and ‘Duchangwudou’ were cloned and inserted into the binary vector pGreenII-0800-LUC (Figure 2A) to evaluate the regulatory effect of GT1CONSENSUS motif (InDel_-904normal) on the activity of *GmRAV* promoter.

**Figure 2.**
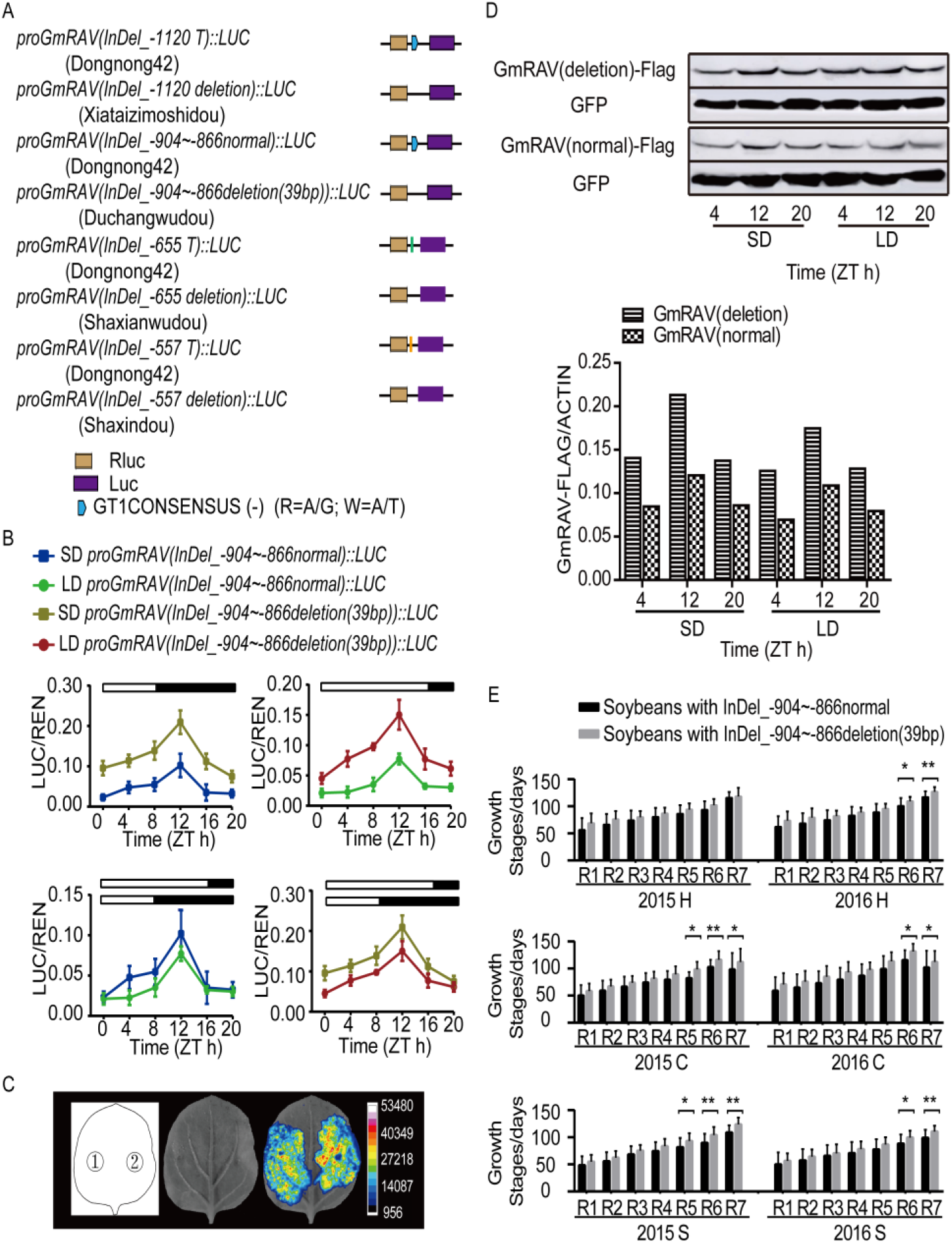
The *GmRAV* promoter activity assay and the *GmRAV* protein expression. (A) Schematic diagram of four pairs of constructs *proGmRAV (InDel_-1120 T)::LUC* and *proGmRAV (InDel_-1120 deletion)::LUC, proGmRAV (InDel_-904~-866 normal)::LUC* and *proGmRAV (InDel_-904~-866 deletion (39bp))::LUC, proGmRAV (InDel_-655 T)::LUC* and *proGmRAV (InDel_-655 deletion)::LUC, proGmRAV (InDel_-557 T)::LUC* and *proGmRAV (InDel_-557 deletion)::LUC*. (B) The LUC/REN relative expression activity of the effector constructs *pro (InDel_-904~-866deletion(39bp))::LUC* and *pro (InDel_-904~-866normal)::LUC* in *Nicotiana benthamiana* in LDs and SDs, respectively. (C) Luciferase activity of *proGmRAV (InDel_-904~-866deletion(39bp))::LUC* and *proGmRAV (InDel_-904~-866normal)::LUC* effector constructs at 12 h after dawn under SDs. 1: *proGmRAV (InDel_-904~-866normal)::LUC*; 2: *proGmRAV (InDel_-904~-866deletion(39bp))::LUC*. (D) The protein expression profiles of GmRAV driven by *proGmRAV (InDel_-904~-866 normal)* promoter and *proGmRAV (InDel_-904~-866 deletion (39bp))* promoter in *Nicotiana benthamiana* in LDs and SDs, respectively. Experiments were performed three times independently. The ratios of signal strength of GmRAV proteins and GFP loading control were calculated. (E) The flowering and mature periods of soybeans with polymorphisms InDel_-904~-866deletion(39bp) or InDel_-904~-866normal in six environments. Student’s t-test, *P < 0.05; **P < 0.01. 2015 H: 2015 Harbin; 2015 C: 2015 Changchun; 2015 S: 2015 Shenyang; 2016 H: 2016 Harbin; 2016 C: 2016 Changchun; 2016 S: 2016 Shenyang.

The analysis on the diurnal expression rhythm of effector constructs *proGmRAV (InDel_-904~-866 deletion(39bp))::LUC* and *proGmRAV (InDel_-904~-866 normal)::LUC* in *N. benthamiana* showed that the construct *proGmRAV (InDel_-904~-866 deletion(39bp))::LUC* showed stronger LUC/REN relative activity than that of *proGmRAV (InDel_-904~-866 normal)::LUC* both in SDs and LDs, which reached a peak around 12 h after dawn and decreased toward dawn (Figure 2B). And the LUC/REN relative activity of *proGmRAV (InDel_-904~-866 deletion(39bp))::LUC* was 2.05-fold (*P*-value<0.01) and 1.95-fold (*P*-value<0.01) more than that of *proGmRAV (InDel_-904~-866 normal)::LUC* in both SDs and LDs at 12h after dawn (Supplemental Figure 3). Similar results at 12h after dawn in SDs were shown through the luminescence signals of the plant imaging system (Figure 2C). Meanwhile, three pairs of constructs *proGmRAV (InDel_-1120 T)::LUC* and *proGmRAV (InDel_-1120 deletion)::LUC*, *proGmRAV (InDel_-655 T)::LUC* and *proGmRAV (InDel_-655 deletion)::LUC*, *proGmRAV (InDel_-557 T)::LUC* and *proGmRAV (InDel_-557 deletion)::LUC* were also constructed (Figure 2A) and transfected to *N. benthamiana*, but the differences among them were not significant at 12h after dawn in SDs and LDs (Supplemental Figure 3).

We also analyzed the diurnal *GmRAV* protein expression rhythm of constructs *proGmRAV(InDel_-904~-866 normal)::GmRAV-3F6H-35S::GFP* and *proGmRAV(InDel_-904~-866 deletion (39bp))::GmRAV-3F6H-35S::GFP* by infiltrating *Agrobacterium* into tobaccos in both LD and SD conditions, respectively. The gene encoding *GmRAV* protein in the two soybean varieties above is the same, but the promoter was different. The *GmRAV* protein level driven by *GmRAV* promoter (InDel_-904~-866 normal) was lower than that driven by *GmRAV* promoter (InDel_-904~-866 deletion (39bp)) at ZT 4h, ZT 12h and ZT 20h in both LDs and SDs (Figure 2D).

Therefore, the difference of *GmRAV* expression pattern between soybeans ‘Dongnong 42’ and ‘Duchangwudou’ might be caused by GT1CONSENSUS motif (InDel_-904normal) that decreased the mRNA transcript and protein level of *GmRAV* in ‘Dongnong 42’ compared with that GT1CONSENSUS motif mutation (InDel_-904deletion(39bp)) in ‘Duchangwudou’. Furthermore, we found that *GmRAV* promoter was regulated by the day length, the LUC/REN relative activities of both *proGmRAV* (InDel_-904~-866deletion(39bp))::LUC and *proGmRAV* (InDel_-904~-866normal)::LUC were generally higher in SDs by comparing with that in LDs, and the differences were significant at 12h after dawn (Figure 2B, Supplemental Figure 4). Also, the analysis on the protein expression patterns of *GmRAV* showed that the accumulation levels of the *GmRAV* protein whether driven by *GmRAV* promoter (InDel_-904~-866 normal) or by *GmRAV* promoter (InDel_-904~-866 deletion (39bp)) in SD conditions were higher than that in LD conditions and reached the peak at ZT12 (Figure 2D), which was consistent with the *proGmRAV*::LUC expression pattern (Figure 2B). The change of GmRAV protein levels was caused by the change of transcriptional level.

Meanwhile, the average maturity of 235 soybeans with InDel_-904normal was earlier than that of 14 soybeans with InDel_-904deletion(39bp) in six environments, respectively (Figure 2E, Table S8). Other growth periods were similar (Figure 2E, Table S8). ‘Dongnong 42’ with less *GmRAV* protein level matured earlier than ‘Duchangwudou’. Therefore, we inferred that GT1CONSENSUS motif in *GmRAV* promoter probably shortened growth periods R5-R7 by reducing *GmRAV* gene expression.

### GmRAV inhibits flowering time and maturity in soybean

To further clarify the biological functions of *GmRAV* gene with promoter (InDel_-904~-866 normal) in soybean, we transformed the *proGmRAV* (InDel_-904~-866 normal)*::GmRAV-3F6H* construct into soybean (Figure 3). In total, three stable T_3_ transgenic lines *GmRAV-ox-5, GmRAV-ox-7* and *GmRAV-ox-14* were obtained (Supplemental Figure 5 A, B). They displayed later R1, R3 and R5, especially significantly later R7 in SDs. Moreover, they also displayed later R1and R3, especially significantly later R5 and R7 in LDs. *GmRAV-ox* soybean plants showed shorter height and fewer numbers of leaves compared with control plants, even some plants grow the first trifoliate leaf from the cotyledon instead of true leaf under SDs and LDs (Figure 3A-C). As expected, T_6_ transgenic *GmRAV-i* soybean displayed earlier flowering time and maturity, and increased plant heights (Figure 3A and 4B) (Zhao et al., 2012). We analyzed the protein expression profiles of GmRAV in *proGmRAV::GmRAV-3F6H* plants in LDs and SDs and found that GmRAV protein driven by the native *GmRAV* promoter was higher in SDs conditions compared to LDs and reached the peak at ZT12 by western blot assay (Figure 5F), which was consistent with the transient expression assays of *GmRAV* promoter activity and GmRAV protein levels in tobaccos (Figure 2B,D). Furthermore, the change of GmRAV protein levels was caused by the change of transcriptional level, in accordance with the *GmRAV* mRNA induction at ZT12 in SDs compared with that in LDs (Lu et al., 2014). The results above confirmed our deduction that higher levels of GmRAV protein expression inhibited flowering and maturity time. *GmRAV* was a growth period repressor which guaranteed both vegetative and reproductive phase long enough to allow necessary energy reserved to be accumulated in SDs and also ensured a strict control of flowering and maturity time.

**Figure 3.**
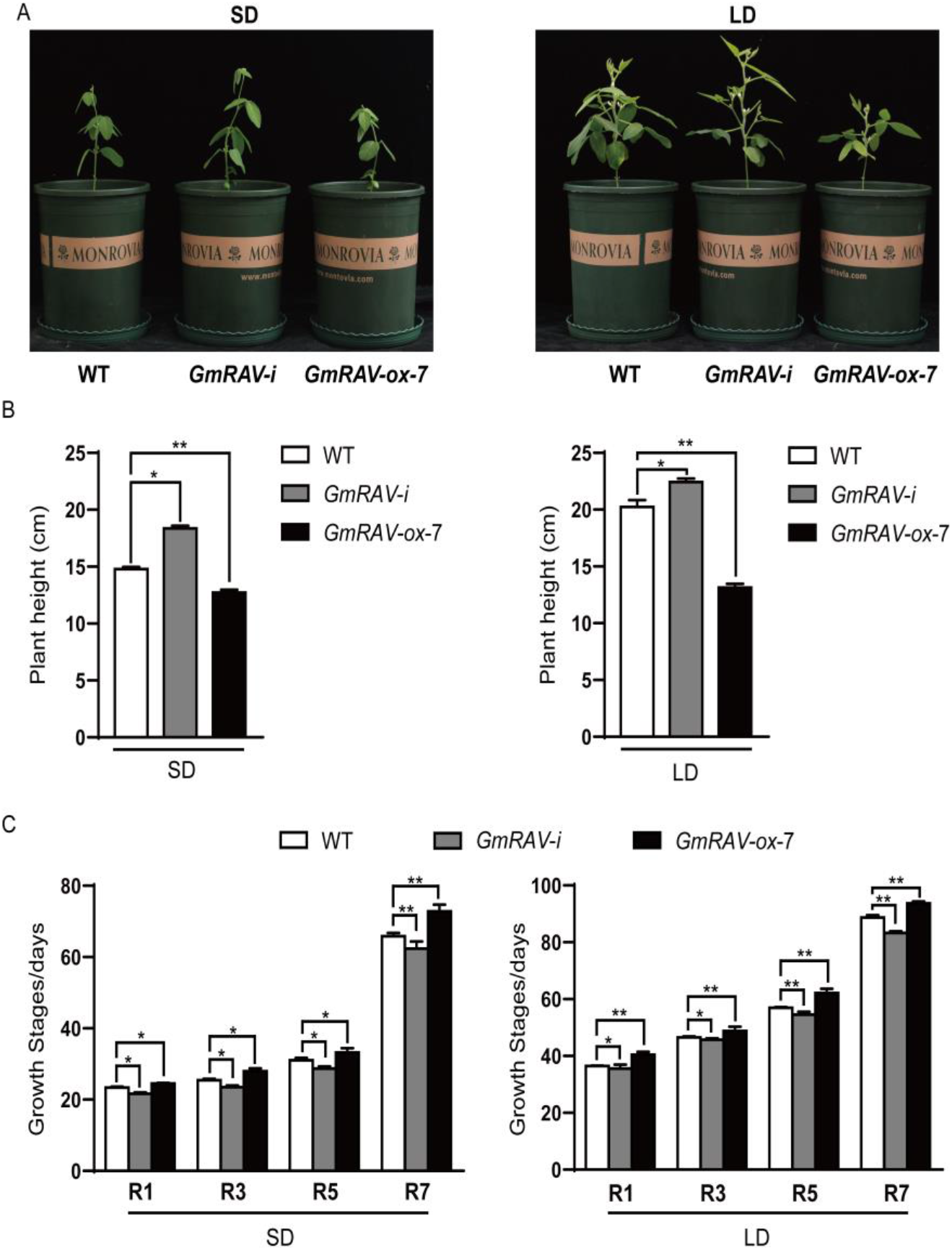
GmRAV inhibit flowering time and maturity. (A) Phenotypes of the T_6_ generation *GmRAV-i-3* and T_3_ generation*GmRAV-ox-7* soybean under LD (16 h light/8 h dark) at 38 d and SD (8 h light/16 h dark) conditions at 25 d after emergence. (B) Flowering time, plant height, yield per plant under LD and SD conditions Data are shown as Means±SEM (n ≥ 15). Student’s t-test, *P < 0.05; **P < 0.01.

### Genome-wide identification of GmRAV-Target Genes by DAP-seq and RNA-seq

To identify the DNA binding sites and the target genes of GmRAV in soybean and further elucidate the potential mechanism by which *GmRAV* inhibited flowering and maturity, we performed DAP-seq and whole transcriptomic RNA sequencing (RNA-seq). Based on DAP-seq data from the two biological repeats, 776 GmRAV-binding sites were identified and grouped into 645 genes, which were referred to as GmRAV-target genes (Supplemental Data Set 1). These binding sites comprised gene bodies and their flanking regulatory sequences, including 2-kb upstream regions that are assumed to harbor promoter regions and 2-kb downstream regions that are assumed to contain terminator regions. Further analysis revealed that 86.34%, 6.83%, and 6.83% of the GmRAV binding sites were located in defined gene bodies, promoter regions and terminator regions, respectively (Figure 4A). According to the consensus sequences at the detected GmRAV-binding sites, putative GmRAV-binding motifs were predicted using the Hypergeometric Optimization of Motif EnRichment (HOMER) software (Heinz et al., 2010). Based on the predictions, and the GmRAV binding motifs [C(A/G/T)A(C)ACAA(G/T)A(C/T)A(G/T)] and [C(T/A)A(C/T) C(T/G)CTG] were identified (Figure 4B), which was consistent with the report that in Arabidopsis, the AP2 and B3 domains of *AtRAV1* protein cooperatively bind to specific target sequences (CAACA and CACCTG motifs)(Kagaya et al., 1999).

**Figure 4.**
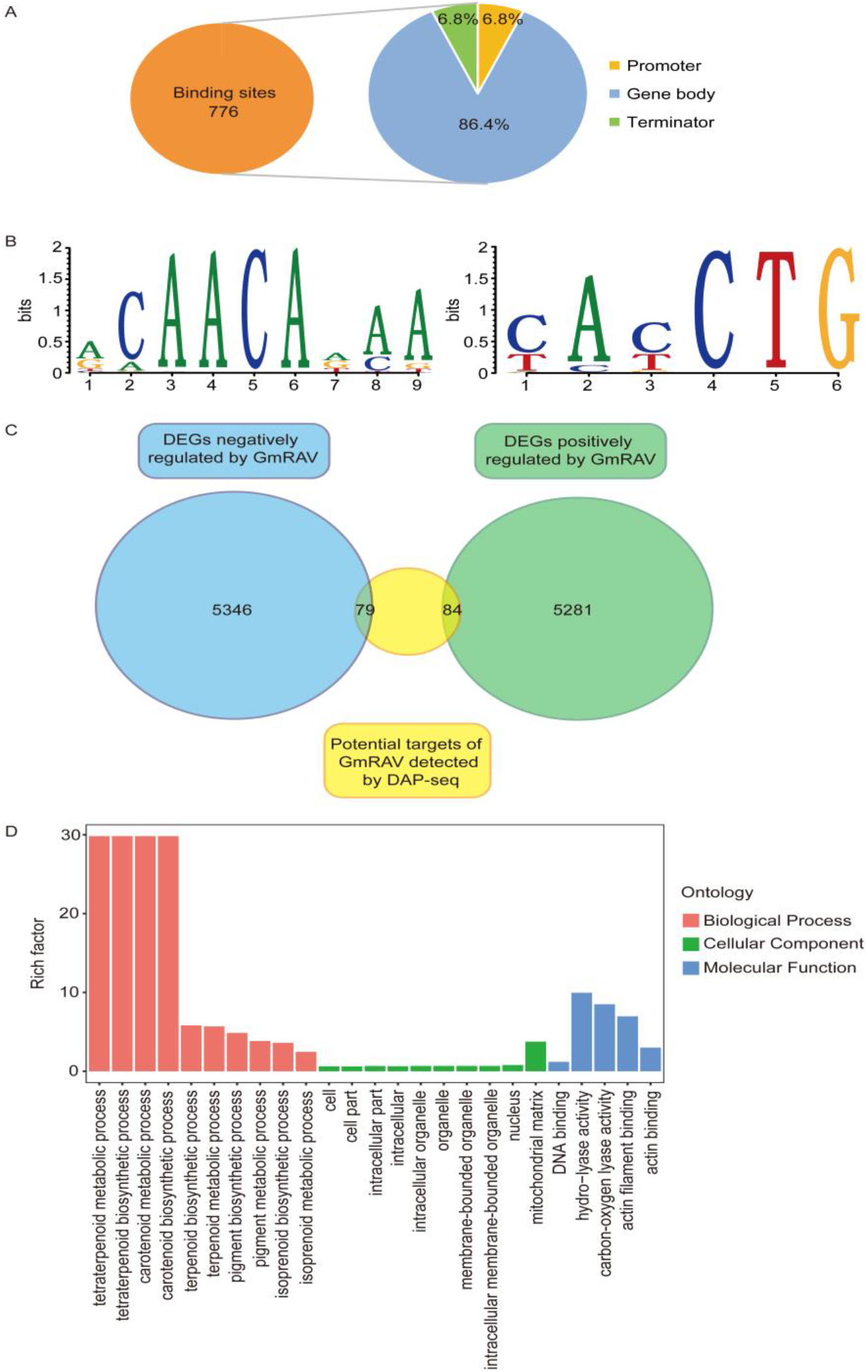
Genome-wide identification of GmRAV-binding sequences and target genes. (A) Distribution of the locations of binding sites relative to target genes. Definitions: promoter is a sequence within 2 kb upstream of transcription start site; terminator is the sequence within 2 kb downstream of the transcription termination site; gene body includes 5’ UTR, CDS, intron, and 3’ UTR. (B) Motif analysis of GmRAV-binding sequences using Homer software. For the distribution of a single motif among the sequences, the parameter was set to zero or one per sequence. (C) Venn diagram showing a comparison between RNA-seq and DAP-seq results. DEGs, differentially expressed genes. (D) Gene ontology (GO) classification for DEGs with GmRAV-binding sites detected.

To establish the regulatory roles of GmRAV, we identified the GmRAV targets and additional genes whose transcriptional activities are associated with relative abundance of the GmRAV transcripts based on RNA-seq analysis. Transcriptomic data from trifoliate leaves from 15-day-old wild-type and *GmRAV-ox-7* transgenic soybean at ZT12 under SDs were generated and compared, and a total of 10627 differentially expressed genes (DEGs) between wild type and the GmRAV-ox transgenic line were detected (Supplemental Data Set 2). These include 5281 genes upregulated and 5346 genes downregulated by GmRAV overexpression (Figure 4C). Enriched GO terms of these differently expressed genes were related to the regulation of reproductive process. Flowering promoters such as *GmFT2a*, *GmFT5a,* two *FRUITFULL (FUL)* homologs *GmFULa (Glyma.06g205800)* and *GmFULb (Glyma.04g159300)* were downregulated and flowering repressor *GmPRR37 (Glyma.12G074400)* was upregulated by *GmRAV* overexpression.

Combining the results from the DAP-seq and RNA-seq analyses, 163 GmRAV-target genes were determined to be putatively directly bound and transcriptionally regulated by GmRAV (Figure 4C; Supplemental Data Set 3). Of these candidates, 84 genes were upregulated and 79 genes were downregulated in *GmRAV-ox-7* (Figure 4C). A gene ontology (GO) analysis of the identified GmRAV-target genes indicated that they were enriched in GO terms related to tetraterpenoid metabolic/biosynthetic process, carotenoid metabolic metabolic/biosynthetic process, DNA binding, implying that *GmRAV* could regulate different biological processes that affect plant growth and development (Figure 4D). Among these, 15 genes whose promoters were also bound by GmRAV in the DAP-seq experiment were identified. Of these, ten genes were upregulated and five genes were downregulated in *GmRAV-ox-7* soybean (Supplemental Data Set 3).

To test whether GmRAV could directly bind to the [C(A/G/T)A(C)ACAA(G/T)A(C/T)A(G/T)] and [C(T/A)A(C/T) C(T/G)CTG] motif, the in vitro binding of GmRAV to the CAACA and CACCTG motifs was analyzed using EMSAs. GST-GmRAV could dramatically reduce the migration of the 35-bp and 40-bp probe, indicating that GmRAV could directly bind to both CAACA and CACCTG motif (Figure 5A). Reciprocal competitive EMSA showed strong and specific binding of GmRAV to the CAACA and CACCTG motifs.

**Figure 5.**
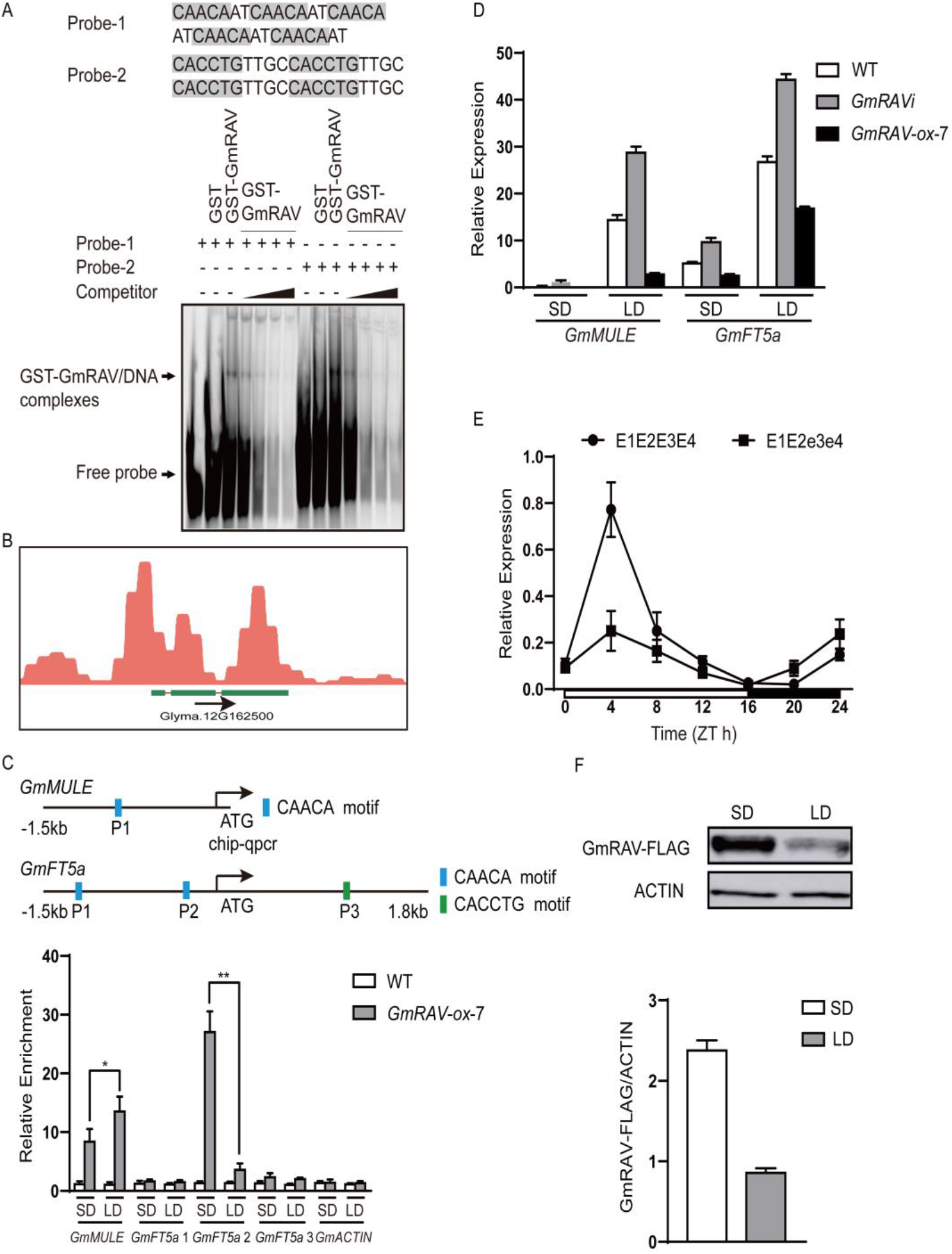
Validation and expression analyses of selected two GmRAV target genes. (A) EMSA assay showing the specific binding of GmRAV to CAACA and CACCTG binding motifs. Competition for GmRAV binding was performed with a cold probe (competitor) to the labeled probe at molar ratios of 1:1, 1:5, 1:10, and 1:25. The CAACA and CACCTG binding motifs are shaded in black. GST served as a negative control. (B) Peak graphs showing the DAP-seq raw reads at the indicated gene loci in Integrative Genomics Viewer. The arrows indicated the directions of transcription, and the green bars indicated the transcript of gene. (C) Binding of GmRAV protein to the *GmMULE* and *GmFT5a* promoters was enhanced by SDs. Relative enrichment of GmRAV-binding fragments from the regulatory region of GmRAV detected by ChIP–qPCR using FLAG-Antibody treatment with *GmRAV-ox-7* seedlings grown in LDs or SDs for 15 days and wild type as a control. Relative enrichment of *GmMULE* fragment containing GCAACATCA motif and *GmFT5a* fragments containing CAACA and CACCTG motifs marked 1 to 3 indicated three regions from the regulatory region of GmRAV detected by ChIP–qPCR. The relative enrichment of the wild type fragment was set as 1.0 and those of other fragments were adjusted accordingly. The *GmACTIN4* locus was used as a negative control. The top image shows the locations of the PCR fragments in *GmFT5a* gene. Values are shown as mean ± standard errors of the mean (SEM) from three biological replicates. Student’s t-test, *P < 0.05; **P < 0.01. Upper panel: physical locations of fragments harboring putative motifs are shown in the schematic diagram. (D) Quantitative real-time PCR analysis of the transcript levels of *GmMULE* and *GmFT5a* in WT, *GmRAV-i-3*, and *GmRAV-ox-7* seedlings grown in LDs or SDs for 15 days. Data are Means±SEM of three biological replicates. (E) Quantitative real-time PCR analysis of the transcript levels of *GmRAV* in NIL *E1E2E3E4* and *E1E2e3e4* in LDs for 30 days. (F) The protein expression level of GmRAV in LDs and SDs. *proGmRAV (InDel_-904~-866 normal)::GmRAV-3F6H* soybean seedlings grown in LDs or SDs for 15 days. Experiments were performed three times independently. Means±SEM were calculated from ratios of signal strength of GmRAV proteins and ACTIN loading control.

### *GmRAV* inhibited the transcriptional function of transcription factor *GmMULE* and *GmFT5a*

We further validated the GmRAV target, *GmMULE (Mutator-like transposable elements)* containing MULE transposase domain *(Glyma.12G162500)* which might be associated with light signal (Figure 5B). Transcription factors *FHY3* and *FAR1* encode Mutator-like transposases to modulate phytochrome A-signaling pathway (Hudson et al., 2003; Jiang et al., 2004). *FHY3* and *FAR1* are transcription factors derived from mutator like transposable element during evolution. However, the functional role of *GmMULE* is not known yet. The loci of *GmMULE* located at promoter regions with strong enrichment of the GCAACATCA motif were selected to further verify the DAP-seq data. The GmRAV target *GmMULE* which might be involved in soybean flowering and maturity, based on the DAP-seq analysis, but the differential expression of flowering promoter *GmFT5a (Glyma.16G044100)* not detected in DAP-seq analysis but significantly downregulated by *GmRAV* overexpression in RNA-seq analysis (Supplemental Data Set 2), which was in accordance with the late-flowering phenotype. Furthermore, *TEM* (*GmRAV* homologue) was mainly expressed in the vascular tissue of leaves and controlled flowering time by repressing the expression of *FT* through direct binding to the CAACA and CACCTG motifs in the *FT* promoter in *Arabidopsis* (Lee et al., 2007). Therefore, *GmFT5a* were also further analyzed to determine whether GmRAV could directly bind them by ChIP-qPCR. The two CACCA motifs were located in promoter and CACCTG was located in the exon of *GmFT5a* in soybean (Figure 5C). The leaves of 15-day-old *proGmRAV::GmRAV-3F6H-ox-7 (GmRAV-ox-7)* transgenic lines grown in LDs or SDs at ZT 12h were used to perform ChIP-qPCR to verify potential GmRAV-binding sites with wild-type sample as a negative control. We harvested the samples at ZT12 in SDs and LDs, corresponding to an abundant GmRAV protein time point (Figure 5F). We found significant binding of GmRAV to the *GmMULE* promoter region and *GmFT5a* (P2) promoter upstream of the ATG site at 230 bp (containing CAACA motif) both in SDs and LDs. We found little binding of GmRAV to the other sites of *GmFT5a* (Figure 5C). When the GmRAV protein level was low in LDs (Figure 5F), light signals significantly increased the binding capacity of GmRAV to the promoter of *GmMULE* to more significantly repress the expression of *GmMULE* in LDs than in SDs, a process that appeared to be distinct from the rising levels of GmRAV protein by ChIP-qPCR of GmRAV. However, we found that the association of GmRAV to the promoters of *GmFT5a* was clearly increased in SDs compared with LDs at ZT12 consistent with the rising levels of GmRAV proteins in SDs (Figure 5C). The short daylength induced increase in GmRAV binding occurred, when the GmRAV protein level was significantly altered (Figure 5C), which indicated that short daylength signals sharply increased the binding capacity of GmRAV to the promoter of *GmFT5a*, consistent with the rising levels of GmRAV proteins. Furthermore, the time course-dependent expression analysis revealed that the mRNA expression level of *GmRAV* in the *E1E2E3E4* NIL was repressed compared with *E1E2e3e4* in LDs. The mRNA expression level of *GmRAV* reached a maximum at 4 h after dawn and decreased to their original levels by dusk in LDs (Figure 5E).

Moreover, the *GmMULE* and *GmFT5a* promoters were bound by GmRAV and whose expression levels in leaves were both elevated in *GmRAV-i* and decreased in *GmRAV-ox-7* soybean plants (Figure 5D). We also tested the functional interaction of GmRAV on CAACA in and *GmMULE* and *GmFT5a* promoter in vivo by using transient expression assay in *N. benthamiana* (Fig. 7A, B). In this system, the LUC reporter gene was driven by *GmMULE* promoter containing a CAACA motif and *GmFT5a* promoter containing two CAACA motifs at C-terminal. When co-infiltrating *Agrobacterium* expressing *proGmRAV(InDel_-904~-866 normal)::GmRAV-3F6H-35S::GFP* or *proGmRAV(InDel_-904~-866 deletion (39bp))::GmRAV-3F6H-35S::GFP* effectors together with the *proGmMULE::LUC* and *proGmFT5a::LUC* reporter into tobacco leaves, irrespectively, both the activity of LUC specially decreased in accordance with the amount of expressed GmRAV-3F6H proteins (Figure 6B,C), thus demonstrating that *GmRAV* could inhibit the transcriptional activation activities of *GmMULE* and *GmFT5a* in both SDs and LDs. Together, our results suggested that GmRAV protein repressed the expression of *GmMULE* and *GmFT5a* under both SD and LD conditions by directly binding to their promoters. GmRAV acted downstream of *E3/E4* to inhibits flowering by repressing *GmFT5a/GmMULE* expression (Figure 8).

**Figure 6.**
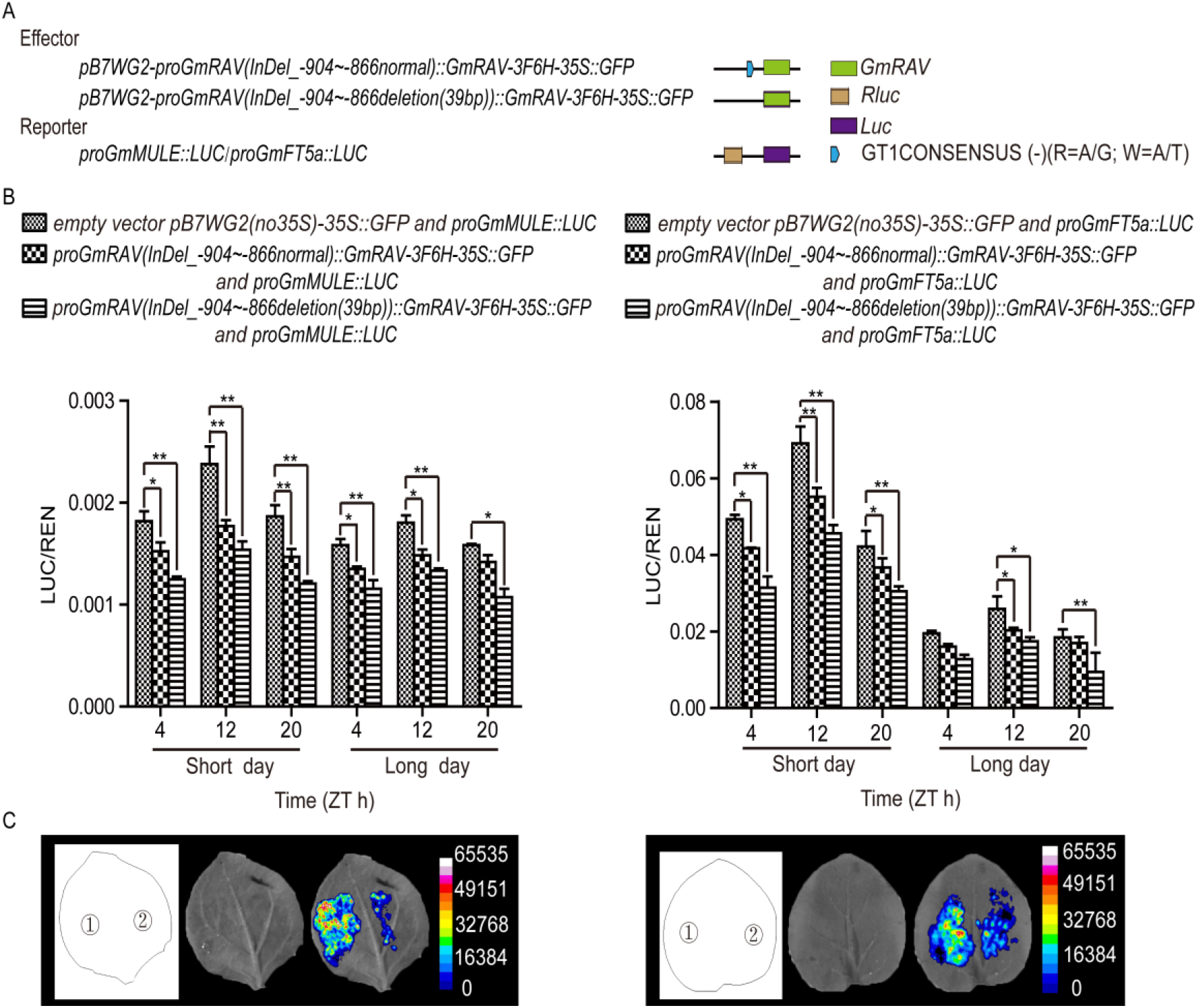
The effect of GmRAV protein on the *GmMULE* and *GmFT5a* promoter activity. (A) Schematic diagram of effector constructs *pB7WG2-proGmRAV (InDel_-904~-866 normal)::GmRAV-3F6H-35S::GFP*, *pB7WG2-proGmRAV (InDel_-904~-866 deletion (39bp))::GmRAV-3F6H-35S::GFP*, empty vector *pB7WG2 (no35S)-35S::GFP* and reporter constructs *proGmMULE::LUC* or *proGmFT5a::LUC*. (B) The effect of GmRAV protein on the *GmMULE* and *GmFT5a* promoter activity. Relative luciferase activity was monitored in tobacco leaves cotransfected with effector and reporter constructs at three time points in LDs and SDs. Quantitative real-time RT-PCR analyses of *GFP* mRNA abundance was calculated to measure the concentration of injection of Agrobacterium containing GmRAV constructs. *GmACTIN4* was used as an internal control. Values are Means±SEM (n=3). Student’s t-test, *P < 0.05; **P < 0.01. The activities of firefly LUC were normalized by the activities of *35S:Renilla LUC*. Results represent Means±SEM of six independent samples. (C) Luciferase assay of *pB7WG2-proGmRAV (InDel_-904~-866 deletion (39bp))::GmRAV-3F6H-35S::GFP*, *pB7WG2 (no35S)-35S::GFP* and *proGmMULE::LUC* or *proGmFT5a::LUC* effector constructs at 12h after dawn in SDs. D-luciferin was used as the substrate of LUC. Left panel 1: *pB7WG2 (no35S)-35S::GFP*+*proGmMULE::LUC;* 2: *pB7WG2-proGmRAV (InDel_-904~-866 deletion (39bp))::GmRAV-3F6H-35S::GFP*+*proGmMULE::LUC*. Right panel 1: *pB7WG2 (no35S)-35S::GFP*+*proGmFT5a::LUC;* 2: *pB7WG2-proGmRAV (InDel_-904~-866 deletion (39bp))::GmRAV-3F6H-35S::GFP*+*proGmFT5a::LUC.*

**Figure 7.**
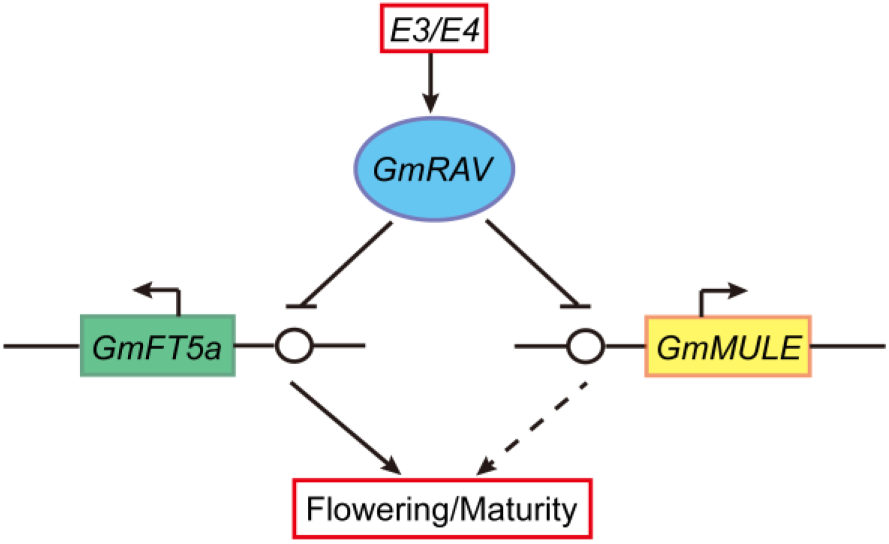
A proposed model showed the roles of GmRAV in flowering time and maturity regulations in soybean. Two phytochrome A photoreceptors *E3*/*E4* induce the GmRAV expression. GmRAV protein directly binds to the promoters of *GmFT5a* and *GmMULE* genes to suppress their transcription to delay flowering and maturity. The solid arrows indicate activation of transcription, dotted lines denote relationship that are not established to be direct, and solid lines ending with a dash denote repression of transcription.

## DISCUSSION

### The natural variation of *GmRAV* contributed to ecological adaption and genetic improvement of soybean

Soybean vegetative and reproductive growth stages like inflorescence development, flowering, pod filling, leaf senescence, and maturity were all influenced by photoperiod. Flowering time is critical for the reproductive success of the plant and is highly variable among natural population. Genetic variations of photoperiod responses enable crops to adapt to various latitudes (Watanabe et al., 2011a). The non-uniform distribution pattern of polymorphisms occurred in the whole *GmRAV* region, the nucleotide diversity of the non-coding region of *GmRAV* was remarkably higher than that of the coding region. As previous study argued for naturally occurring promoter variants create functional diversity, there was an adaptive advantage of one allele of *FT* over the other in particular growth conditions (Liu et al., 2014), thus, we speculated the variation of *GmRAV* promoter contributed adaptability to different environments and requirements in the evolution of development. The deletion of a single nucleotide in *J* was associated with soybean growth adaptation in tropical and low altitude regions (Yue et al., 2017). The natural variations of *GmRAV* related to growth periods such as InDel_-904normal>deletion(39bp), InDel_-557T>deletion, SNP_-293C>A, InDel_-201~-201TA>deletion exhibited geographic distribution patterns consistent with the maturity characteristics, the soybeans with minor polymorphisms InDel_-904deletion(39bp), InDel_-557deletion, SNP_-293A, InDel_-201~-201deletion flowered and matured later, promoting adaptation to lower latitudes. This could provide an important theoretical basis for ecological adaption and genetic improvement of soybean in modern molecular breeding.

### GT1CONSENSUS motif regulated the levels of *GmRAV* expression by photoperiod response

Natural variation in *GmPRR37* affected photoperiodic flowering and contribute to regional adaptation of soybean (Koo et al., 2013). GT1CONSENSUS motif was involved in light-responsive actions (Xue et al., 2018). We also found that the natural variation InDel_-904~-866normal>deletion(39bp) causing a mutation in cis-acting element GT1CONSENSUS motif of *GmRAV* promoter could extremely significantly delayed soybean growth periods, by increasing the protein level of *GmRAV* in soybean ‘Dongnong 42’ (early maturity) compared with that in soybean ‘Duchangwudou’ (late maturity) in SDs and LDs. The cis-acting element GT1CONSENSUS motif (InDel_-904~-866normal) of *GmRAV* promoter controlled the growth period and shortened R5-R7 by reducing the expression level of *GmRAV* in soybean. Consistently, *GmRAV-ox* and *GmRAV-i* transgenic plants displayed more significant changes of R5-R7 in SDs or LDs.

### GmRAV binds to two cis-acting elements to regulate gene transcription

RAV proteins contain two DNA-binding domains, an AP2/ERF and a B3 DNA-binding domain. The CAACA and CACCTG sequences have previously been identified as the DNA recognition sites of RAV1 (Song et al., 2017). TEM as a repressor could directly bind to 5’ UTR region of *FT* containing CAACA and CACCTG to repress *FT* expression in Arabidopsis (Castillejo and Pelaz, 2008). Identifying GmRAV binding motifs will help elucidate the function of RAV transcription factors in soybean. DAP-seq was performed to identify the DNA binding sites of GmRAV in soybean and elucidate the potential flowering mechanism. Two GmRAV binding motifs were identified, which was consistent with the report that in Arabidopsis, the AP2 and B3 domains of *AtRAV1* protein cooperatively bind to specific target sequences (CAACA and CACCTG motifs). In vitro binding of GmRAV to CAACA and CACCTG motifs and functional interactions with CAACA in the promoter were further verified by electrophoretic mobility shift assays (EMSAs). Interestingly, soybean GmRAV could bind just a motif such as CAACA motif of the promoter to repress the expression of downstream *GmMULE* and *GmFT5a* gene. Members of the Mutator DNA transposon superfamily are termed ‘MULEs’ (Mutator-like elements) (Le et al., 2000). In plant genomes there are large numbers of non-autonomous Mutator elements known collectively as ‘Pack-MULEs’, that have lost their transposase genes and instead carry fragments of host genes (Jiang et al., 2004). MULE-derived genes act as transcription factors that modulate the light response in Arabidopsis. Transcription factors *FHY3* and *FAR1* encode Mutator-like transposases to modulate phytochrome A-signaling pathway(Hudson et al., 2003; Jiang et al., 2004). We predicted that *GmMULE* might be also involved in light response, there still needed to validate its function in soybean. The ChIP data of GmRAV biding *GmFT5a* presented here also implicated binding were more enriched in SDs than in LDs, and short daylength signals sharply increased the binding capacity of GmRAV to the promoter of *GmFT5a*, consistent with the rising levels of GmRAV proteins by inhibiting *GmFT5a* expression.

### GmRAV acting downstream of *E3/E4* delayed soybean growth period by repressing *GmFT5a* transcriptional activity

*FT* homologs in soybean belong to the phosphatidylethanolamine binding protein (PEBP) family and serve as major points of integration in flowering time control. Many *FT* genes are expressed in leaves under flower-inducing conditions. FT proteins are thought to be transported from leaves to shoots or lateral apical meristems through the phloem, where they induce the development of floral meristems(Golembeski and Imaizumi, 2015). *GmFT5a* was strongly induced under SDs, caused early flowering when overexpressed (Kong et al., 2010), and *ft5a* mutants delayed flowering (Cai et al., 2020). *E3E4* and *GmFT4* repressed flowering(Zhai et al., 2014). *FT* levels are the result of a quantitative balance between the respective promoter and repressive activities. The photoreceprtor *E3(PHYA3)/E4(PHYA4)* repressed the mRNA level of *GmRAV(Glyma.10G204400)*, so we deduced that GmRAV acted downstream of *E3/E4* to inhibits flowering by repressing *GmFT5a/GmMULE* expression.

The mechanism of RAV action seemed to be similar in the both Arabidopsis and soybean species. To avoid precocious flowering, Arabidopsis TEMs directly target the florigens *FT (Castillejo and Pelaz, 2008)*, whereas GmRAV also repressed *GmFT5a* directly. Upregulation of GmRAV expression was necessary to provide the plant with the competence to respond to inductive short-day photoperiods, as it reduced *GmFT5a* accumulation to avoid earlier flowering in SDs. The proposal that flowering was a process controlled by a quantitative balance between flower-inducing and -inhibiting substances arose from classical physiological studies on the photoperiodic response. The existence of floral transition repressors possessed a great importance for plant development as they guarantee a vegetative phase long enough to allow necessary energy reserves to be accumulated (Boss et al., 2004) and also ensure a strict control of flowering time. *GmRAV* was a floral transition repressor in soybean which guaranteed a vegetative phase long enough to allow necessary energy reserved to be accumulated in SDs and also ensured a strict control of flowering time. Supporting the functional conservation, the ectopic expression of *OsRAV9* in Arabidopsis plants correlated with the repression of the *TEM* targets *FT* delayed flowering time.

## METHODS

### Plant materials, growth conditions and records of data

Nucleotide variation in non-coding and coding regions of *GmRAV* gene was evaluated in an experimental population consisting of 248 soybean varieties that originated from 22 provinces (latitude 20°13’N to 50°15’N) of China and one (Number 19) originated from Russia (Han et al., 2015) (Table S1). In addition, 30 maturity groups (MGs) standard varieties (Number 250-279) from North American (Singh and Hymowitz, 1999; Sneller, 1994) were also planted for identifying the MG classification of 249 soybean varieties (Table S1). All 279 soybean varieties were planted in experimental farms (Harbin, Changchun, Shenyang) of three different latitudes (45°75’N, 126°63’E; 43°88’N, 125°35’E; 41°44’N, 123°30’E) in China in 2015 and 2016 and grew under natural conditions. Sowing and harvesting have been done in early May and October annually. The field experiment was designed by means of completely randomized block design with two replications, with 2 m length of rows, 0.05m plant spacing, and 0.6 m row spacing. The soybean vegetative stage VE (emergence) and the reproductive stages (R1, R2, R3, R4, R5, R6 and R7) were investigated using the method described by Fehr and Caviness (Fehr and Caviness, 1977). The dates of VE (Vegetative) to R1 (Beginning bloom), R2 (Full bloom), R3 (Beginning pod), R4 (Full pod), R5 (Beginning seed), R6 (Full seed), R7 (Beginning maturity) stages were calculated as phenotypic data. The maturity group classification for 249 soybean varieties was identified compared with 30 North America standard varieties (MG000-MGVIII) (Boerma et al., 2004). Soybean (*Glycine max* ‘Harosoy’ [L58-266; *E1E2E3E4*]) and (OT89-5; *E1E2e3e4*) were grown under LDs (16h/8h light/dark) at a consistent temperature of 25°C.

### Sequencing and polymorphism analysis

Genomic DNA of 249 varieties (except for MGs) were extracted by CTAB extraction procedure (Attitalla, 2011). The full-length sequences consist of 1,459-bp promoter region, 457-bp 5’ untranslatable region (5’UTR) and 1,056-bp coding sequence (CDS) of *GmRAV* were amplified by overlapping polymerase chain reaction (PCR) with three pairs of *GmRAV-1*-F *and GmRAV-1*-R*, GmRAV-2*-F and *GmRAV-2*-R, *GmRAV-3*-F and *GmRAV-3*-R primers, respectively (Table S2). The PCR products were purified and sequenced by the Sanger DNA Sequencing Methods (Sanger et al., 1977). Multiple sequence alignment and polymorphism detection were conducted according to a reference genome sequence from Williams 82 using Clustal X 1.83 software (Kim and Joo, 2010). The nucleotide diversity and Tajima’s D neutrality test were calculated by DnaSP.5.10 (Librado and Rozas, 2009; Tajima, 1989). Selective pressure of *GmRAV* in 249 soybean germplasm resources was estimated under three distinct models, FEL, SLAC and MEME (Delport et al., 2010). LD (linkage disequilibrium) decay of full-length sequences of *GmRAV* in 249 soybean varieties was analyzed by TASSEL Version 5.0 (Bradbury et al., 2007).

### Polymorphism-trait association and statistical analysis

Association study was conducted by TASSEL Version 5.0 based on Mixed Linear Model (MLM) with 1000 permutations (Bradbury et al., 2007). The 249 soybean varieties had been sequenced by specific locus amplified fragment sequencing (SLAF-seq) in our lab, the genetic data was taken as covariance of association analysis (Han et al., 2015; Lipka et al., 2012). Only polymorphisms with MAF greater than 4% were used for association analysis on account of mutations with low frequency without the ability to identify the relevance to phenotypes (Korte and Farlow, 2013). The association analysis results were plotted by the Snp.plotter package in R environment (Luna and Nicodemus, 2007). Significant or extremely significant correlations between polymorphisms and growth periods were determined by Bonferroni thresholds -log_10_ (*P*-value)≥1.3 or -log_10_ (*P*-value)≥2.0.

### Plasmid Construction

To generate four pairs of constructs of *GmRAV* promoter driven LUC, including *proGmRAV (InDel_-1120 T)::LUC, proGmRAV (InDel_-1120 deletion)::LUC, proGmRAV (InDel_-904~-866 normal)::LUC, proGmRAV (InDel_-904~-866 deletion (39bp))::LUC, proGmRAV (InDel_-655 T)::LUC, proGmRAV (InDel_-655 deletion)::LUC, proGmRAV (InDel_-557 T)::LUC, proGmRAV (InDel_-557 deletion)::LUC*, the native *GmRAV* promoters with polymorphisms InDel_-1120 T, InDel_-1120 deletion, InDel_-904~-866 normal, InDel_-904~-866 deletion (39bp), InDel_-655 T, InDel_-655 deletion, InDel_-557 T, and InDel_-557 deletion were amplified from genomic DNA of ‘Dongnong 42’, ‘Xiataizimoshidou’, ‘Dongnong 42’, ‘Duchangwudou’, ‘Dongnong 42’, ‘Xiataizimoshidou’, ‘Dongnong 42’, and ‘Xiataizimoshidou’ using forward primers *proGmRAV::LUC-1120-*F, *proGmRAV::LUC-904*-F, *proGmRAV::LUC-557/655*-F, *proGmRAV::LUC-557/655*-F and reverse primer *proGmRAV::LUC*-R (Table S2), respectively. To generate *GmMULE*, *GmFT5a* promoters driven *LUC* constructs *pGmMULE::LUC, pGmFT5a::LUC*, the promoter DNA was amplified from genomic DNA of ‘Dongnong 42’ using *GmMULE::LUC*-F and *GmMULE::LUC*-R, *pGmFT5a::LUC*-F and *pGmFT5a::LUC*-R primers (Table S2), respectively. The PCR products were purified and cloned into binary vector pGreenII-0800-LUC linearized by *Sma*I using In-Fusion cloning system (Clontech, USA), respectively. To generate *proGmRAV::GmRAV-3F6H-35S::GFP* construct to be used for agroinfection in *N. benthamiana*, both *proGmRAV (InDel_-904~-866 normal)::GmRAV-3F6H-pB7WG2-35S::GFP* and *proGmRAV (InDel_-904~-866 deletion (39bp))::GmRAV-3F6H-pB7WG2-35S::GFP* plasmids were constructed. For the FLAG and HIS tag construct, we first synthesized the tandem repeats of 3×FLAG and 6×Histidine (3F6H) tags with *Not*I at 5’ end and *Xba*I at 3’ end (5’-GCGGCCGCCCTGGAGCTCGGTACCCGGG(*Sma*I)GATCCCAGGATCT**GAT TACAAGGATCATGATGGTGATTACAAGGATCACGACATCGACTACAAG GATGACGATGACAAGCACCATCATCACCACCATTGA**TCTCTAGA-3’, the sequences encoding 3F6H tag were in bold) (Song et al., 2012). The synthesized products above were cloned into *Not*I-*Xba*I sites of *pENTRY* vector (named *pENTRY-3F6H*) which contained the 3F6H sequence in the C terminus of the cloning site, and the sequences were verified. *GmRAV* promoter and full-length coding region of *GmRAV* were amplified from genomic DNA of ‘Dongnong 42’ and ‘Duchangwudou’ using *proGmRAV::GmRAV-3F6H*-F and *proGmRAV::GmRAV-3F6H*-R primers (Table S2), respectively. The PCR products were purified and cloned into *pENTRY-3F6H* vector linearized by *Sma*I using In-Fusion cloning system to construct recombinant vector *proGmRAV::GmRAV-3F6H-pENTRY*. To generate *pB7WG2* vector without 35S expression cassette (*pB7WG2-no35S*), *pB7WG2* plasmid was digested by *Spe*I and *Sac*I to remove 35S promoter and the cohesive ends of the linear fragment were blunted by T4 DNA Polymerase. The blunt ends were ligated by T4 DNA ligase. LR reaction was conducted by *proGmRAV::GmRAV-3F6H-pENTRY* and *pB7WG2-no35S* to generate *proGmRAV::GmRAV-3F6H-pB7WG2* fusion expression vector. Both the *proGmRAV::GmRAV-3F6H-pB7WG2* fusion expression vector and the plant GFP expression vector pCAMBIA1302 were double digested by HindIII and SmaI, and then the digested target fragment was inserted into *35S::GFP-pCAMBIA1302* vector to produce *proGmRAV::GmRAV-3F6H-35S::GFP vector*. The constructs above were used for agroinfection in *N. benthamiana*. For the cloning of *GmRAV* into the *pDEST15* vector to express protein in *E. coli*, the full-length coding region of *GmRAV* was amplified by PCR using *GmRAV-TOPO*-F and *GmRAV-TOPO*-R primers (Table S2) and cloned into *pENTR/D-TOPO* (Life technologies) and then transferred to the expression vector *pDEST15* vector through LR reaction to generate the *GST-GmRAV* fusion vector. The construct was transformed into the *E. coli* competent cell line *BL21 (DE3)* (Transgene, Beijing, China) to produce the recombinant proteins.

### Transient assay of *GmRAV* promoter and three promoters affected by *GmRAV* protein in *N. benthamiana*

The recombinant constructs were introduced into *Agrobacterium GV3101* and subsequently transformed into *N. benthamiana* (Sheikh et al., 2014). The transient activity of four pairs of recombinant vectors *proGmRAV (InDel_-1120 T)::LUC, proGmRAV (InDel_-1120 deletion)::LUC, proGmRAV (InDel_-904~-866 normal)::LUC and proGmRAV (InDel_-904~-866 deletion (39bp))::LUC, proGmRAV (InDel_-655 T)::LUC, proGmRAV (InDel_-655 deletion)::LUC, proGmRAV (InDel_-557 T)::LUC, proGmRAV (InDel_-557 deletion)::LUC* were assayed using dual luciferase assay kit (Promega, USA) and multifunctional microplate reader TECAN Infinite 200 PRO (Tecan Schweiz, Männedorf, Switzerland).

The constructs *proGmRAV(InDel_-904~-866 normal)::GmRAV-3F6H-35S::GFP, proGmRAV(InDel_-904~-866 deletion (39bp))::GmRAV-3F6H-35S::GFP* and *pGmMULE::LUC, pGmFT5a::LUC* were simultaneously transferred into *N. benthamiana* to measure transient assay of the promoters affected by *GmRAV* protein using the same measurement method described above, respectively. Three independent experiments were performed and each experiment was repeated three times to obtain reproducible results. The luminescence signal was captured using Amersham Imager 600 (General Electric Company: https://www.ge.com/) after spraying 1 mM luciferin (Heliosense: http://www.heliosense.com/) on *N. benthamiana* leaves.

### Transformation of soybean cotyledon nodes

The construct *proGmRAV::GmRAV-3F6H-pB7WG2* were transferred into the *A. tumefaciens* strain, *LBA4404*. Transgenic soybean ‘Dongnong 50’ plants expressing the *proGmRAV::GmRAV-3F6H-pB7WG2* construct were produced as described previously (Zhao et al., 2018).

### Immunoblot analysis

To measure the protein expression of *GmRAV* gene driven by *GmRAV* promoter (InDel_-904~-866 normal) and *GmRAV* promoter (InDel_-904~-866 deletion (39bp) after agroinfection in *N. benthamiana*, and the protein expression of *GmRAV* gene driven by *GmRAV* promoter (InDel_-904~-866 normal) in transgenic soybean, whole protein extract was extracted from frozen-ground leaves in the extraction buffer [150 mM NaCl, 50 mM Tris (pH 7.5), 10% glycerol, 5 mM EDTA, 0.5% Triton X-100, 0.5% (SDS), 1 mM DTT, 2 mM Na_3_VO_4_, 2 mM NaF and EDTA-free protease inhibitor tablet (Pierce)]. 20μg protein in each samples were separated by 10% SDS-polyacrylamide gel electrophoresis gels, and transferred to nitrocellulose membranes, and were detected using a HRP-conjugated anti-FLAG antibody (A8592, Sigma). GFP protein was detected by HRP-conjugated anti-GFP antibody (ab6663, Abcam) in tobacco, whereas ACTIN protein was detected by mouse anti-actin monoclonal antibody (MAB1501, Chemicon international), followed by a HRP-conjugated goat anti-mouse IgG (H+L) Secondary Antibody (G-21040, Thermo Fisher Scientific) in soybean. GFP and Actin was used for normalization of a protein in whole extract in tobacco and soybean, respectively. Super Signal West Pico Chemiluminescent substrate kits (Thermo Fisher Scientific) and the signal was detected by chemiluminescence imaging (Amersham Imager 600). All experiments were performed at least three times with independent biological replicates.

### DAP-Seq and data processing

DAP-seq was carried out as previously described with minor modifications (O’Malley et al., 2016). gDNA was extracted from young soybean leaves, fragmented, purified using AMPure XP beads (Beckman Coulter, Inc., Indianoplis, IN, USA). Libraries were constructed using the NEXTFLEX Rapid DNA-Seq Kit (PerkinElmer, Inc., Austin, TX, USA) according to the instruction manual. The coding sequence of GmRAV was cloned into a pFN19K HaloTag T7 SP6 Flexi expression vector. Halo–GmRAV fusion protein was expressed using the TNT SP6 Coupled Wheat Germ Extract System (Promega) following the manufacturer’s specifications for expression. Expressed proteins were directly captured using Magne Halo Tag Beads (Promega). The protein-bound beads were incubated with adaptor-ligated gDNA fragments, washed and eluted. Eluted DNA fragments were PCR amplified and sequenced on an Illumina NavoSeq. Reads were mapped to the phytozome genome for *Glycine max* (Wm82.a2.v1). DAP-seq peaks were called using Macs2 (Zhang et al., 2008). Association of DAP-seq peaks located upstream or downstream of the transcription start site (TSS) within 2 kb were analyzed using Homer (Heinz et al., 2010), based on the General Feature Format (GFF) files. Motif discovery was performed using GEM (Genome Wide Binding Event Finding and Motif Discovery) (Guo et al., 2012) to identify the motifs to which GmRAV bound.

### RNA-seq, statistical analysis and qRT-PCR validation of differentially expressed genes

The methods used for RNA-seq were described previously (Zhao et al., 2018). For RNA-seq, trifoliate leaves from 15-day-old wild-type and *GmRAV-ox-7* transgenic soybean at ZT12 under SDs were harvested for RNA expression analysis. The cDNA library preparation, RNA-seq sequencing, and assembly were performed on the Illumina Nova 6000 sequencing platform (SeqHealth Tech Co., Ltd Wuhan, China). For identification of differentially expressed genes, log_2_ ratios were calculated with the reads per kilobase of exon model per million mapped reads (RPKM) value of every gene with P-value ≤0.05 and fold change≥2 were divided into upregulated and downregulated transcripts. GO annotation analyses were completed using the BLAST2GO software (https://www.blast2go.com/), an automated tool for the assignment of GO terms.

### Electrophoretic mobility shift assay (EMSA)

Oligonucleotide probes containing (CAACA) and (CACCTG) motifs were synthesized and labeled with Cy5 at 5’ ends, respectively. Probe sequences were shown in Supporting Information Table S2. For competition with unlabeled probe, unlabeled probe was added to the reactions. A total of 1 μg of cell extracts containing GST-GmRAV protein were incubated with 20 μM of Cy5-labeled probe in a binding buffer [25 mM Hepes·KOH, pH 7.5, 40 mM KCl, 0.1 mM EDTA, 5% (vol/vol) glycerol, 2.5 mM DTT, 0.2 μg·μL^−1^ BSA, 15 ng poly dI-dC] and appropriate amounts of unlabeled competitor DNA (1-,5-, 10- and 25-fold molar excess with respect to the labeled probe). After incubation for 30 min at room temperature, samples were separated by electrophoresis on native polyacrylamide gels (8% acrylamide and bis-acrylamide, 39:1). Fluorescent gel images were obtained by using a Typhoon FLA 9500 Biomolecular Imager (GE Healthcare Life Sciences).

### ChIP-qPCR

Wild-type and soybean transgenic *proGmRAV::GmRAV-3F6H-ox-7* lines were grown for 15 days in LDs or SDs in growth chamber at 25°C. Approximately 1 g of trifoliate leaves from wild-type and transgenic cucumber lines were harvested at ZT 12h, and fixed with 1% (v/v) formaldehyde for 15 min under vacuum infiltration. Final concentration of 0.125M glycine was subsequently added to quench the cross-linking reaction. After grinding the seedlings, nuclei were isolated and lysed, and the chromatin solution was then sonicated to approximately 200-1000 bp DNA fragments. Immunoprecipitation reactions were performed using anti-FLAG antibody (Monoclonal ANTI-FLAG® M2 antibody produced in mouse, F1804, Sigma-Aldrich). The complex of chromatin antibody was captured with protein G beads (Invitrogen), and DNA was purified using a QIAquick PCR purification kit (QIAGEN). The enrichment of DNA sequence segments in gene promoters was chosen to perform qPCR. Three biological repeats and three technical replicates were performed for each sequence segment. *GmACTIN* was used as the internal gene control. The primer pairs used in ChIP-qPCR are listed in (Table S2).

ACCESSION NUMBERS Sequence data from this article were submitted to the National Center for Biotechnology Sequence Read Archive (NCBI-SRA) database under the BioProject no. PRJNA682336 and accession nos. SRR13197374, SRR13197373 and SRR13197372 for the DAP-seq data. BioProject no. PRJNA681087 and accession nos. SRR13155194, SRR13155193, SRR13155192, SRR13155191, SRR13155190 and SRR13155189 are for the RNA-seq data.

## SUPPLEMENTAL INFORMATION

Supplemental Information is available at Molecular Plant Online.

## AUTHOR CONTRIBUTIONS

Y.L. and J.S. performed the expression analysis and dual-luciferase assay; L.D. and J.W. performed DAP-seq; J.S. performed soybean transformation and the phenotype observations; M.L. performed SNP detection; C.X. performed EMSA; W.Y. and J.S. performed the gene cloning, soybean transformation, the transcriptome analyses and ChIP qPCR; X.Z., D.L, Y.Z. and J.S. performed the data analysis; L.Z., W.Y., and W.L. wrote the manuscript.

## ACKNOWLEDGMENTS

We thank Dr. Diqiu Yu (Xishuangbanna Tropical Botanical Garden, Chinese Academy of Sciences) and Dr. Fengning Xiang (Shan Dong University) for sharing plasmid materials.

